# Gut microbiome activity contributes to individual variation in glycemic response in adults

**DOI:** 10.1101/641019

**Authors:** Hal Tily, Ally Perlina, Eric Patridge, Stephanie Gline, Matvey Genkin, Vishakh Gopu, Haely Lindau, Alisson Sjue, Iordan Slavov, Niels Klitgord, Momchilo Vuyisich, Helen Messier, Guruduth Banavar

**Author notes:** This study was performed while all authors were at Viome Inc.

## Abstract

Limiting post-meal glycemic response is an important factor in reducing the risk of chronic metabolic diseases, and contributes to significant health benefits in people with elevated levels of blood sugar. In this study, we collected gut microbiome activity (i.e., metatranscriptomic) data and measured the glycemic responses of 550 adults who consumed more than 30,000 meals from omnivore or vegetarian/gluten-free diets. We demonstrate that gut microbiome activity makes a statistically significant contribution to individual variation in glycemic response, in addition to anthropometric factors and the nutritional composition of foods. We describe predictive models (multilevel mixed-effects regression and gradient boosting machine) of variation in glycemic response among individuals ingesting the same foods. We introduce functional features aggregated from microbial activity data as candidates for association with mechanisms of glycemic control. In summary, we demonstrate for the first time that metatranscriptomic activity of the gut microbiome is correlated with glycemic response among adults.

## Introduction

From a public health perspective, preventing elevated levels of blood glucose is a crucial part of mitigating the current epidemic of metabolic diseases including obesity, type 2 diabetes, hypertension, cardiovascular and liver diseases. 9.4% of the US population is diabetic and 26% is prediabetic, creating a large disease burden with associated healthcare costs [CDC 2017]. Daily food choices play the largest role in determining overall blood glucose levels and thus risk for various diseases ([Gutierrez 1998]; [Livesey 2008]; [Jenkins 1985]; [Ludwig 2018]). Tools facilitating the mass adoption of dietary choices to maintain normal glycemic levels would be an important step towards halting the hyperglycemia epidemic.

Popular nutritional understanding largely focuses on food characteristics alone, such as caloric and carbohydrate content. However, there is increasing evidence that glycemic response to the same foods differs significantly among individuals. Recent studies ([Zeevi 2015]; [Mendes-Soares 2019]) have shown that postprandial glycemic response (PPGR) is not only driven by the glycemic index of food, but also the individuals’ phenotypic and molecular characteristics, including the gut microbiome which may have a role in energy metabolism and the regulation of insulin response [Suez 2016]. These studies evaluated postprandial glycemic response (PPGR) in the context of specific populations (Israeli and US midwestern), a small number of standardized meals, and 16S or metagenomic data from the gut microbiome.

In this paper, we present data to demonstrate that the glycemic response to a range of foods varies based on individual differences including gut microbiome *activity*, i.e., metatranscriptomics of the gut microbiome, and anthropometrics. The study presented here generalizes previous results to a significantly larger set of standardized meals (104 unique pre-designed meals) coming from two distinct diet types – omnivore and vegetarian/gluten-free. With the goal of being readily interpretable, this paper provides a concise statistical explanation of the relationship of nutrients, phenotypes, and gut microbiome activity with PPGR, through a multilevel mixed-effects regression model. We also present a gradient boosting machine model that has been optimized for predictive accuracy. Furthermore, we identify multiple significant *functional* microbiome features related to prediction of postprandial glycemic response, indicating that properties correlated with the microbiome affect the processing of carbs as well as leading to an overall difference in baseline blood sugar.

The primary goal of this study was to determine the impact of microbial gene expression (at the functional level) on glycemic response. The most commonly used gut microbiome analysis method, the 16S rRNA gene sequencing, provides poor taxonomic resolution, typically genera that contain many strains with very diverse gene content [Knight 2018]. Metagenomic methods cannot identify some microorganisms (e.g. RNA viruses) and can only predict gene expression based on the gene content, which can be highly erroneous [Bervoets 2019]. We therefore used metatranscriptomics [Hatch 2019], which sequences RNA molecules and provides the primary sequence and read counts for each transcript, allowing us to use the data for quantitative strain-level taxonomic classification and functional pathway analysis ([Gosalbes 2011]; [Bashiardes 2016]; [He 2010]). Due to the challenges posed by RNA instability, the necessity for removal of diverse ribosomal RNAs in stool samples, and complex bioinformatic analyses, metatranscriptomic methods have not been widely used in clinical studies. To our knowledge, this is the first study to demonstrate the application of gut metatranscriptomics in a population-scale dietary study.

## Methods

As shown in Figure 1, we recruited 550 adults (66% female), and tracked their food intake, sleep, activity, and glycemic response for up to 2 weeks. 400 participants were Caucasian, and of the remaining 150 participants, 37% were Asian, 33% were Hispanic, and 30% were Black or Other. The study’s Research Protocol was approved by an accredited IRB committee and all study participants consented to participating in the study. All study participants were at least 18 years old. We obtained a stool sample at participation enrollment, as well as a comprehensive questionnaire describing their lifestyle, preferences, and health history. We collected blood glucose measurements every 15 minutes using a Continuous Glucose Monitor (CGM) sensor that measures glucose levels within the range of 40 to 500 mg/dL [Abbott 2016].

**Figure 1:**
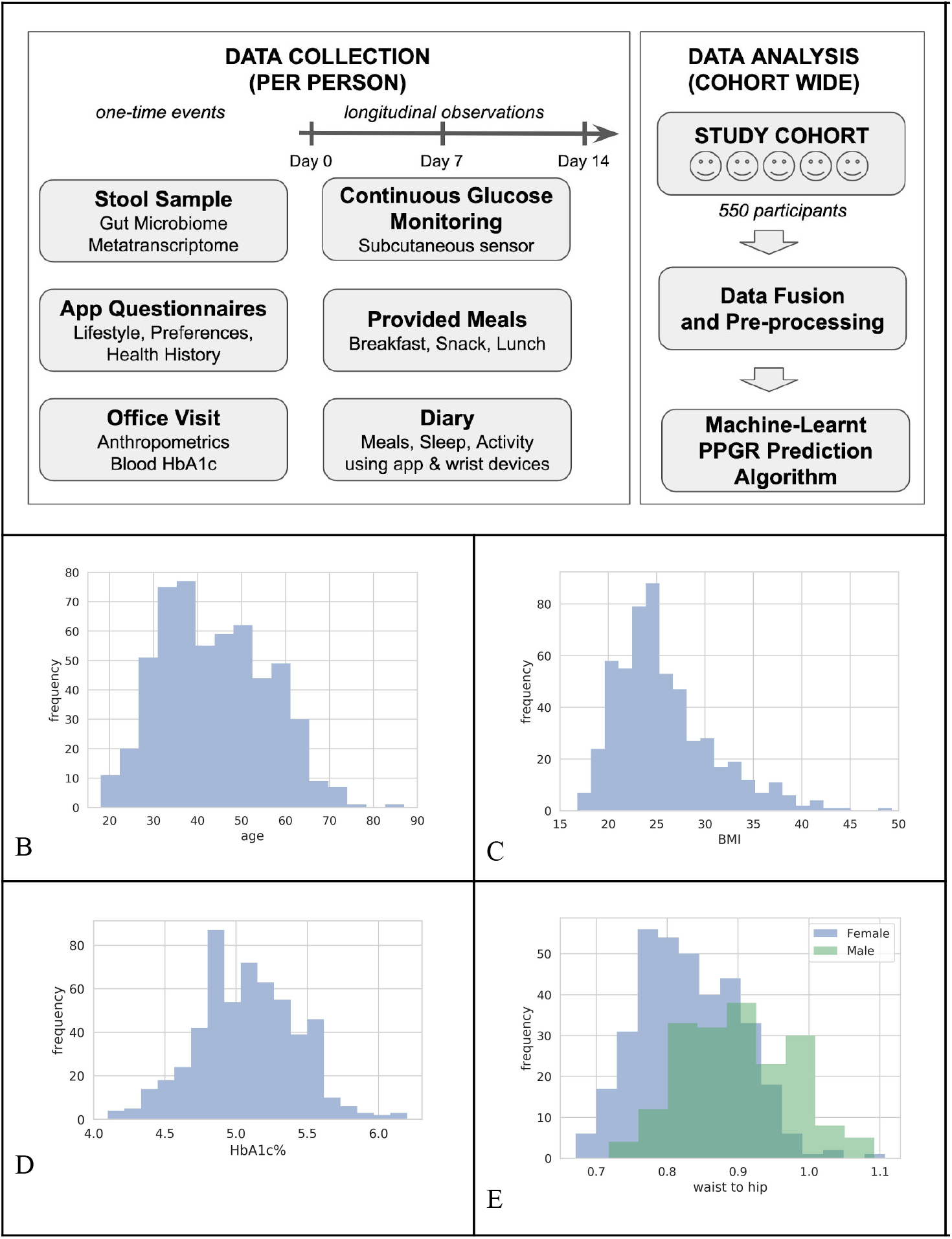
Study Design and Population Characteristics. **1A**. The study cohort had 550 adult participants (66% female). Each study participant provided a stool sample, filled out questionnaires, and made an office visit. Then over 14 days, participants consumed pre-designed meals that were provided, they monitored their blood glucose response, and they kept a diary of their meals, sleep, & activity. At the end of the study, all the data streams were fused, pre-processed, and analyzed as described in this paper. The following exclusion criteria were used: age<18; dietary restrictions that would prevent adherence to any of the study diets; antibiotic use 1 month prior to or during study; skin disease (e.g. contact dermatitis) that precludes proper attachment of the CGM; pregnancy; active neoplastic disease; active neuropsychiatric disorder; myocardial infarction or cerebrovascular accident in the 6 months prior; pre-diagnosed type I or type II diabetes mellitus; HbA1c >= 6.5; or unwilling / incapable of following instructions. **1B**. Age distribution with mean of 43.8 years (SD 12.115). **1C**. 28% of the study population had BMI > 25 and 18% had BMI > 30. **1D**. 4% of the study population were pre-diabetic with HbA1c% > 5.7. **1E**. Waist-to-hip ratio distribution with mean of 0.901 (SD 0.076) for men and 0.832 (SD 0.071) for women.

### Study Meals

As described in Figure 2, participants were provided pre-designed breakfasts, snacks, and lunches (“provided meals”) over 14 days (“day 0” to “day 13”). After lunch, participants were allowed to eat whatever they wanted (“free meals”) without further guidance on the composition, and day 0 consisted of only free meals. Provided meals accounted for 66% of all meals and free meals 34%.

**Figure 2:**
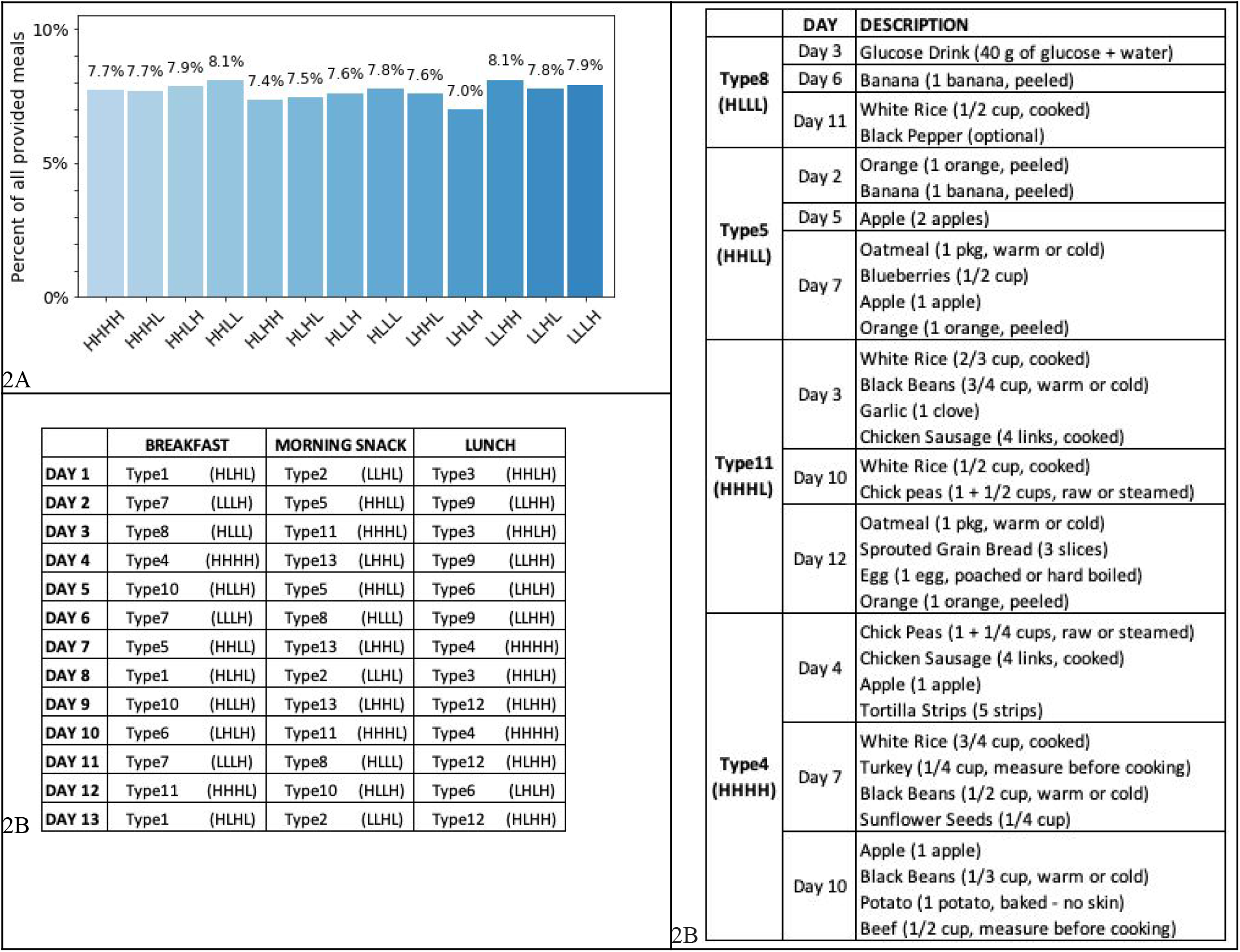
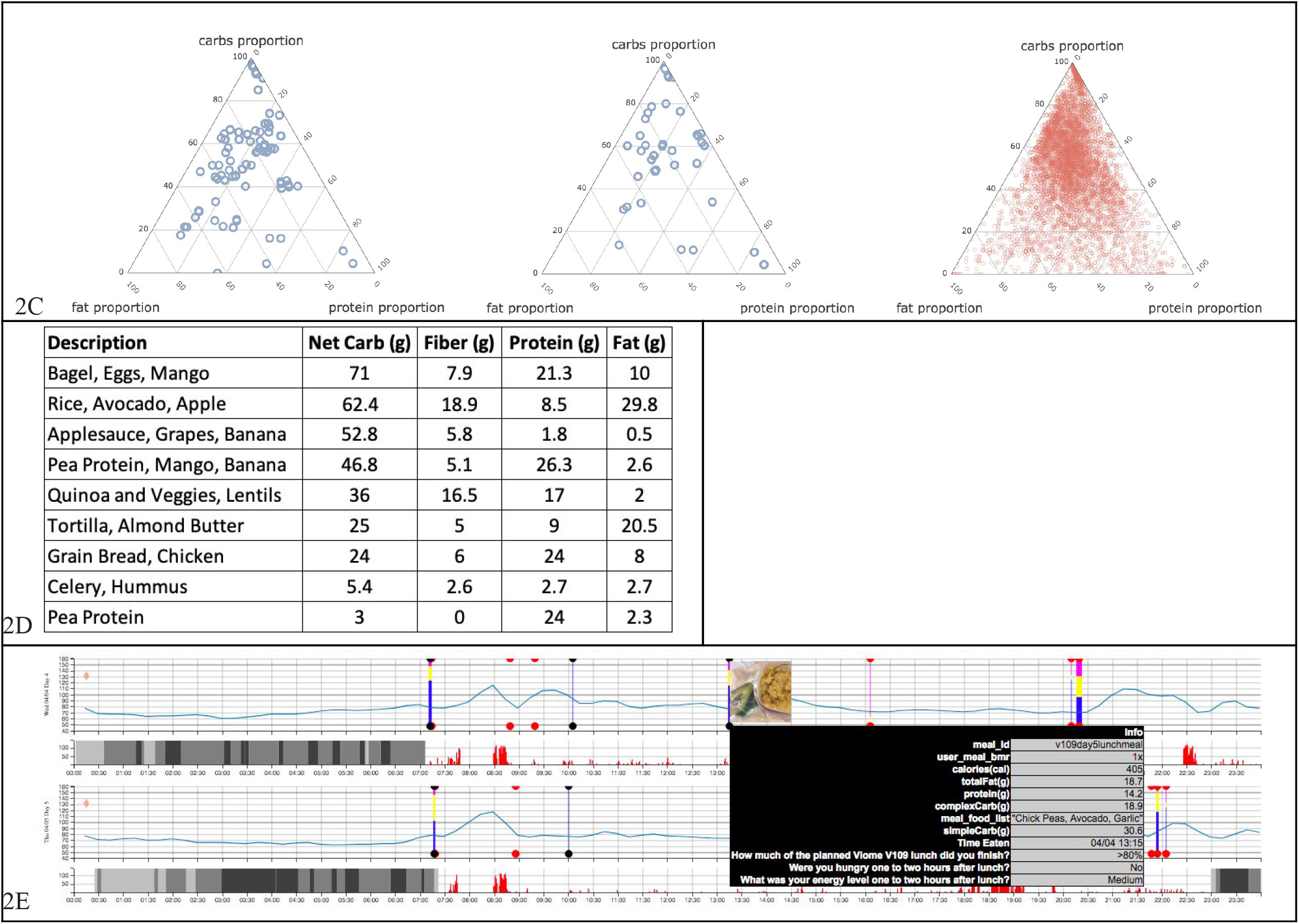
Study meal design and content. **2A**. Percentage of each meal category within all 18,000+ provided meals. Categories were defined using proportions of Carbs (H/L), Fiber (H/L), Proteins (H/L), and Fats (H/L). **2B**. In this omnivore diet example, meals are shown for several of the provided macronutrient groups (carbs - fiber - protein - fat). Meals with similar composition “types” were distributed across the schedule, and schedule days 1 and 8 were repeated meal days. **2C**. Macronutrient proportions of all 71 unique pre-designed omnivore meals (top, blue), all 33 unique pre-designed vegetarian/gluten-free meals (middle, blue), and all free meals (bottom, red). Provided meals account for 66% of all meals, and free meals 34%. **2D**. Macronutrient information for the 9 “repeat meals” which were consumed twice by study participants. Actual servings were adjusted based on the participant’s basal metabolic rate (BMR). **2E**. Data collected for a single participant over 2 days (out of 14). Each row is a single day. The blue curve is the CGM reading collected every 15 mins. Vertical bars are meal events, showing carbs proportion (blue), fat proportion (yellow), and protein proportion (pink). The user interface also visualizes a picture of the meal and the nutrient details. Grey bars represent light and deep sleep. The red histogram next to sleep bars is the tracked physical activity.

Both provided and free meals were recorded by all participants during the entire study period, using a smartphone app (Bitesnap). We obtained macronutrient and micronutrient information from the smartphone app platform for further analysis. Provided meals were pre-loaded into the app. Free meals were loaded by users through selection of custom dishes, ingredients, and quantities.

We asked the participants to not perform intense exercise 2.5 hours before or 2.5 hours after meals, to not start probiotics or prebiotics, to not take vitamins or supplements during the study (with a specific detail to avoid interfering substances as defined in [Abbott 2016]), to not take over-the-counter medication, and to inform the study coordinator if they are prescribed antibiotics during the study.

In order to test our methodology across a range of diets and to support a range of participant preferences, we provided two diet types – omnivore and vegetarian/gluten-free – in different phases of the study. We provided 104 unique pre-designed meals — 71 unique pre-designed meals in the omnivore diet in two separate waves, and 33 unique pre-designed meals in the vegetarian/gluten-free diet in one wave. 140 participants signed up for the omnivore diet and 410 signed up for the vegetarian/gluten-free diet.

To distribute macronutrients across all the meals, we designed the provided meals using a high (H) / low (L) determination of each of the 4 macronutrients – carbs, fiber, protein, and fat – as shown in Figure 2A. In this paper, we use the terms *carbs* and *carbohydrates* interchangeably to mean net carbohydrates, i.e. excluding fiber. High (H) and Low (L) were thresholded based on the proportion of each macronutrient and their daily recommended allowance. For example, day 1 breakfast was high (H), low (L), high (H), low (L), respectively, in carbs, fiber, protein, and fat. An example meal plan is shown in Figure 2B; over the 14 day period (day 0 consisted of only free meals), participants were provided with 39 meals, including one glucose drink. The distribution of macronutrients (Figure 2C) shows the coverage of the provided meals within the space of macronutrient proportions.

Figure 2B includes 3 distinct meals that were repeated on days 1 and 8. A total of 9 distinct meals shown in Figure 2D were repeated over the course of the 14 day diet schedules (omnivore and vegetarian/gluten-free) to collect information regarding intra-person variability; each repeat meal was consumed twice by study participants. By design, these repeat meals were generally high in just one or two macronutrients.

Figure 2E shows the data from all study sources for a single participant over two days, fused into a single visualization. This visualization helped the study administrators to visually inspect the data and ensure that data was properly captured and lined up. Based on this visual inspection, we observed that some of the CGMs malfunctioned with consistent lack of signal, or in some cases the smartphone app meal captured events were out of sync with the glucose curves, which indicated that the participant did not capture the data as instructed (at the point of meal consumption). In either of these cases, we discarded the respective meal data.

Meal data was pre-processed as follows. After discarding meals that were clearly from malfunctioning CGMs or from erroneous data capture, we ended up with 27630 total meals, with 18208 provided meals and 9422 free meals. All provided meals were at least 2.5 hours apart (participants were instructed). Free meals that were within 30 mins were merged, and those within 90 minutes of each other discarded. After all of this pre-processing, each participant provided an average of approximately 50 meals.

### Microbiome Features

Stool samples from participants were processed using our metatranscriptomic method [Hatch 2019] to yield raw microbiome data features including KEGG gene orthologs (or “KOs”) [KEGG] and microbial taxonomy. Our cloud-based bioinformatic pipeline performs read QC/trimming, host read filtering, and taxonomic classification at three taxonomic ranks (strain, species and genus) through sequence alignment to a custom database of more than 110,000 genomes. Functional assignments (KOs) are obtained through alignment to the IGC [Li 2014] and KEGG databases. For the samples provided by the 550 participants in this study, 6587 unique microbial KOs were detected, a mean of 2941.9 per sample (s.d. 541.9); and 1047 unique species were detected, with a mean of 122.7 (s.d. 40.5).

Collections of these raw microbiome data features were aggregated into custom microbiome scores designed to capture the collective functional characteristics as described in the literature. For example, the score *microbiome balance*, is an aggregate assessment of overall ratios of active beneficial and harmful microbes, as well as some diversity metrics. This score is binary with a value of “Low” or “Normal”. All microbiome scores are generated by taking expression data as input, and applying an expert-designed scoring algorithm developed at Viome [Perlina *et al*, in prep.] to derive an overall activity level.

Metabolic and signaling pathway activities are scored using expression levels of genes encoding specific protein functions (KEGG mappings are used primarily), compared with a reference cohort of samples supplied by Viome customers. Scores measure the quantity and expression levels of specific KEGG gene orthologs (KOs) selected due to their specific directional enzymatic roles, pathway topology, or significance in the functional literature. The more key genes expressed, and the higher their expression levels, the higher the resulting score.

## Results

### Meal Effects

For the remainder of our analyses we look at *incremental area under the curve* (iAUC), a standard assessment of PPGR ([Cheng 2018]; [Wolever 1986]). We define this as the integrated area under the CGM curve over 120 minutes, relative to the baseline CGM measured at the time of the meal, i.e., the difference between the area above baseline and area below baseline. Note that this measurement can be negative due to decline in blood sugar level below the baseline over time, especially after activity, and due to noise from the measurement device. The actual iAUC values for all meals, the repeated meals from Figure 2D, and the glucose drink are shown in Figure 3A.

**Figure 3:**
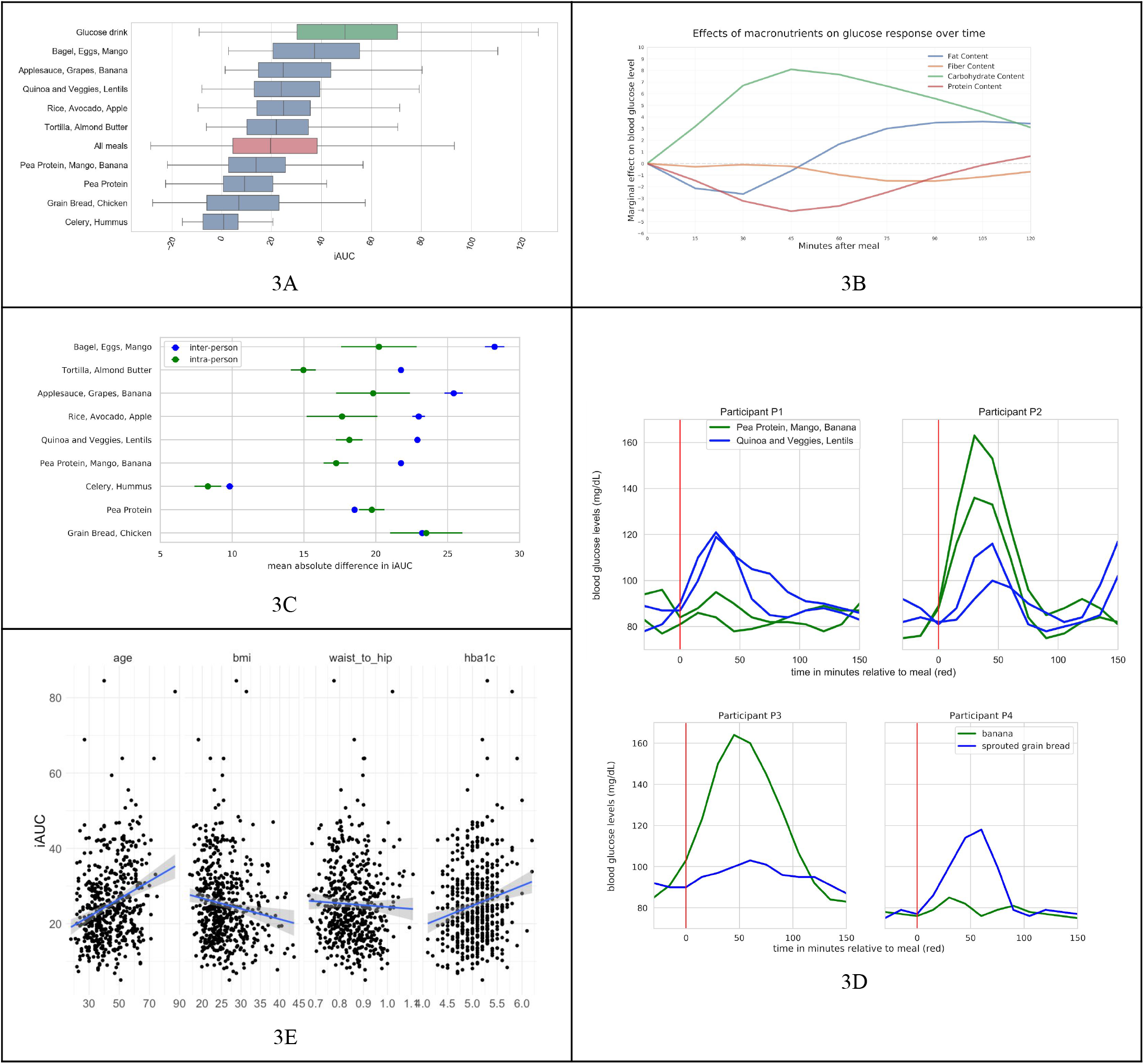
Overview of postprandial glycemic response data. **3A**. iAUC values (in mg / dl-h) for all meals, provided and free (red bar); the 9 repeated meals from Figure 2D, (blue bars); and the glucose drink (green bar). Boxplots show the interquartile range; whiskers cover the middle 95th percentile. **3B**. Marginal effect of macronutrients on glucose response over time, across all meals, across all participants. Each timepoint is a linear regression of iAUC on all four standardized macronutrients. **3C**. Inter- and intra-person variability for 9 repeated provided meals. X axis is the mean absolute difference in iAUC. Points indicate the mean absolute difference in response between two consumptions of the meal by one person (green), and mean absolute difference in response between all pairs of different people (blue). Bars indicate standard error. Y axis is in descending order of difference between inter-person and intra-person variability. **3D**. Examples of individual variation in glycemic response. Ingestion of two of the repeated meals (blue and green lines) result in opposite blood glucose response in two participants (top). Ingestion of two free meals results in opposite blood glucose response in two participants (bottom). **3E**. Relationships between anthropometric characteristics and per-participant average iAUC across all meals (provided free).

Figure 3B was constructed after modeling iAUC with standardized macronutrient values (i.e., z-scores). Linear regression of iAUC on all four of the standardized macronutrients was performed at each timepoint after the meal. The plot shows the learned weights for each of the standardized macronutrients at each of these timepoints, and this reveals the magnitude and time-course of the macronutrient effects. Meals with more carbohydrates led to increased postprandial glucose (PPG), peaking 45 minutes after the meal, while meals with more fiber led to a diminished and delayed PPG. Protein and fat suppress and delay the response as shown.

Figure 3C illustrates the variability in responses to a single meal, for all meals that were repeated within the provided diet (Figure 2D). Intra-person variability is the difference in a participant’s response to a single meal when eaten on two occasions.

### Predictive Model Development and Evaluation

In this section we first present a linear *multilevel mixed-effects* or *hierarchical* model [Gelman 2007] of PPGRs based on the data described above. The linear model allows us to provide a concise description of the relationships between nutrients, anthropometrics, microbiome activity, and PPGR. Additionally, it allows us to derive significance statistics testing the hypotheses that each predictor is relevant in the determination of the PPGR.

Importantly, the inclusion of *random effects* captures individual variation in PPGR due to unobserved factors (unknown properties of the individual, meal, or measurement devices) that may affect the outcome. Including random effects is essential in order to derive conservative hypothesis tests of both the relevance and magnitude of our fixed effects, especially in a repeated-measures design where each person and each meal provide many data points. Our experimental design calls for a multilevel model because of this repetition; each PPGR observation is at a lower level nested within one person and within one meal (higher levels), in a *crossed* or *fully factorial* experimental design (see Figure 4A). For example, without the multilevel model we could not conclusively test the importance of any variable that is constant across all data taken from one person, such as microbiome features.

**Figure 4.**
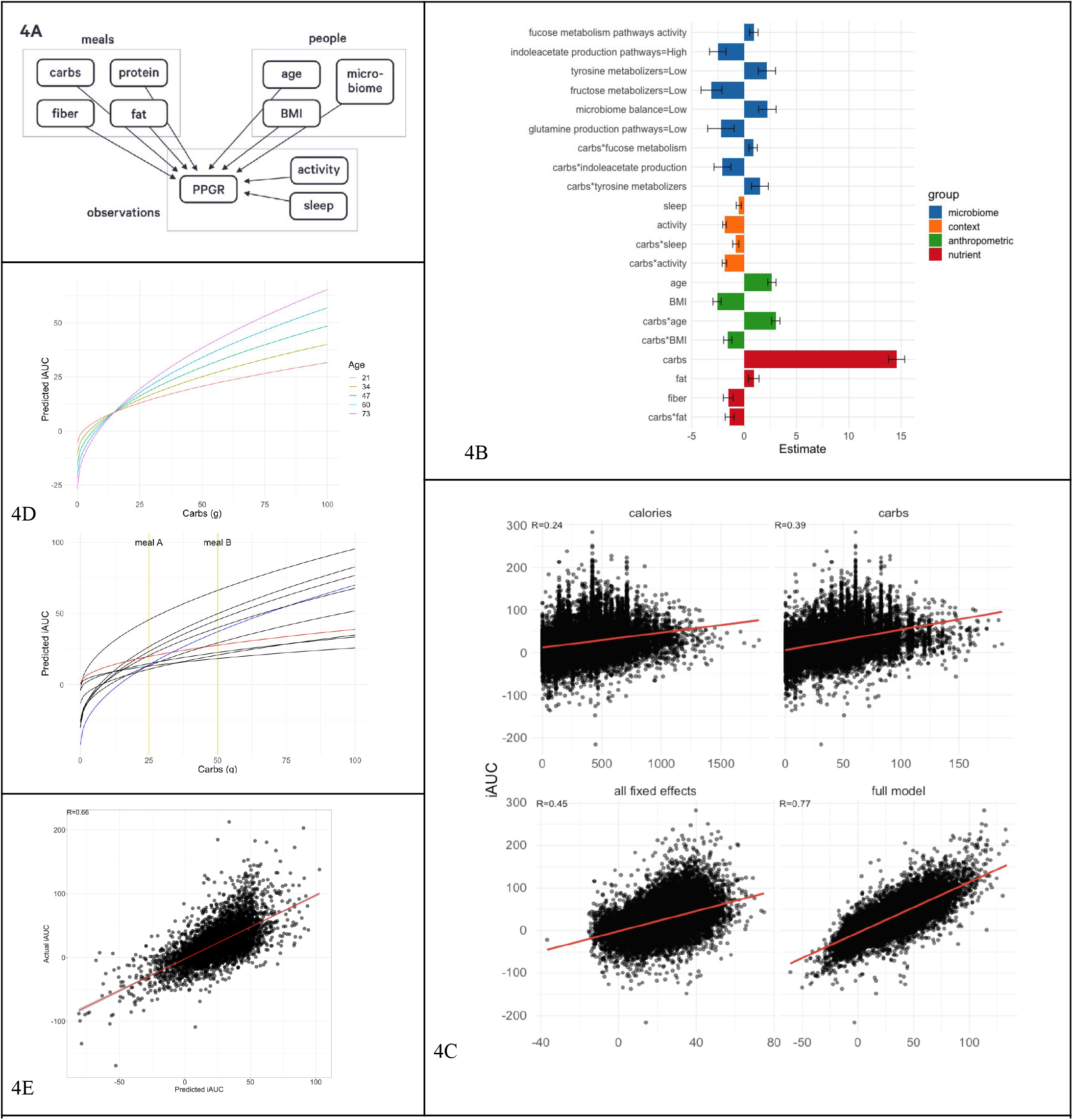
Models for predicting PPGR. **4A**. Nesting of predictors in our repeated-measures crossed experimental design. Plates (boxes) indicate repetition: e.g., there is one measurement of carbs, protein, fiber, fat for each meal. Arrows indicate the possibility of dependence: here, PPGR is estimated as a function of all other variables. **4B**. Fixed effect estimates for all predictors included in the final model. Continuous predictors are standardized (mean is zero, units are standard deviations), meaning the expected response changes by the value of the coefficient when the predictor changes by one standard deviation. All microbiome scores except *fucose* are binary scores, meaning the coefficient is the difference in expected response between the two levels of the score. Error bars show standard error of the estimate. **4C**: Actual glycemic response (iAUC) against (a) calories, (b) carbs, (c) predictions from the model including all fixed effects (standard linear regression), and (d) the fit of the full mixed-effects model. **4D**. (left) Model predictions as a function of carbs and age, holding all other predictors at the baseline. (right) Model predictions as a function of carbs for 10 randomly sampled people, taking into account all person-level fixed and random effects and holding all other nutrient and context variables at the baseline. Two users are highlighted (red and blue) to draw attention to the flip in their predicted response to the two example meals A & B annotated in yellow. **4E**: Actual vs predicted iAUC for a gradient boosting machine (GBM) model. Performance shown is on the best test fold across 5 random splits, with data from 82% of the users used to train and 18% held out for evaluation.

Model development was performed using the *lme4* package in R [Bates 2015]. The model was built incrementally. In the first pass, we determined appropriate transformations for the nutritional and anthropometric variables by visualizing the relationship between each predictor and the iAUC. The effect of carbohydrates appeared to be well-described by the square root transformation, while other predictors were left untransformed. We then fitted a model of these nutritional and anthropometric predictors as fixed effects, and random intercepts and slopes allowing the nutritional effects to vary between people and the anthropometric effects to vary between meals. Likelihood ratio tests showed no evidence for any random slopes of anthropometrics by meal (i.e., responses to all meals are similarly influenced by the participant’s age, BMI, etc).

As the largest effects were associated with carbs, in the second pass we introduced interactions between that and all other nutrients as fixed effects, and with random slopes by participant. Based on visual inspection of their relationship with iAUC, we also introduced fixed effects of two response-level “context” measurements, *minutes of activity during the 2 hours following the meal* and *minutes of sleep during the 3 hours before the meal*. We removed effects not significant by likelihood ratio test at *p<*.*05*.

In the third pass, we introduced fixed effects of our microbiome features, a number of expert-designed scores measuring the activity of crucial metabolic pathways and functions (see discussion section below). We determined a candidate set of scores to add to the model by predicting participants’ average iAUC from each score using t-test (binary) or regression (continuous). 15 scores showed a significant association taken alone and were tested in the full model. After controlling for all other predictors, 6 remained significant or marginally significant.

The final model fixed-effect coefficients are displayed in Figure 4B. Positive coefficients indicate a greater predicted glycemic response. All predictors were significant by likelihood-ratio test at *p<*.*05*, except for glutamine production pathways (*p=*.*08*) and the interaction between tyrosine metabolizers and carbs (*p=*.*06*) which are kept in this model as suggestive, and do not substantially affect the estimation of other coefficients.

Several of the significant predictors were microbiome scores. One of the scores, named microbiome balance, is an aggregate assessment of microbiome balance quantifying overall beneficial and harmful activities (based on the literature), as well as some diversity metrics. This score, when “Low,” showed a negative association with glycemic response. Two other scores are quantified pathway activities representative of overall activity levels of a given set of microbial pathways.

The fucose metabolism pathways activity score considers expression levels of all the genes that encode enzymes known to carry out biochemical reactions that result in processing and catabolic conversions of fucose – a glycan that microbes may obtain from the host’s gut lining mucosal layer, or food components. This quantitative score showed a positive association with glycemic response.

The indoleacetate production pathways activity score considers expression levels of all the genes that encode enzymes known to carry out biochemical reactions that result in production of indole-acetate (or indole acetic acid, IAA). This binary score, when “High,” showed a negative association with glycemic response.

The marginally significant score of glutamine production pathways implies higher glycemic response when “Low.” The score is derived with the same approach as the indoleacetate score, and assess the levels of activity of various pathway axes leading to microbial production of glutamate (or glutamic acid).

Microbial scores for tyrosine and fructose metabolizers are based on functional groups of active microbes known to metabolize tyrosine and fructose, respectively. Tyrosine metabolizers, when “Low,” are directly related to elevated PPGR. Fructose metabolizers, when “Low,” show an inverse relationship to glycemic response.

Figure 4C compares different approaches to predicting iAUC. The first two plots show single predictor models (calories or carbs). We also present predictions from the fixed-effect part of our model after zeroing out certain components. Using all nutrient predictors achieves a similar fit to using the square root of carbs alone (both R=.41, not pictured). Including the microbiome features yields a small but significant improvement in fit (R=.42, not pictured), and adding the other fixed effect predictors (age, BMI, sleep, activity, microbiome) improves the fit further (R=.45, Fig. 4C bottom-left). Finally, the full model including random effects (best linear unbiased predictions) fits the data very well (R=.77, Fig. 4C bottom-right).

Figure 4D (left) shows the influence of age on the modelled relationship between carbs and glycemic response. Figure 4D (right) illustrates the extent of modeled individual differences in this relationship, taking into account all predictors. Two hypothetical meals A and B (similar to repeat meals from Figure 2D with carbs approximately 25g and 50g respectively) are shown to illustrate that two users (blue and red lines) have the opposite glycemic responses (iAUC of 14 & 20 vs iAUC of 37 & 27 respectively) due to the crossover of their iAUC response lines between the two meals — this effect was shown in Figure 3D using the raw data.

### A model optimized for prediction accuracy over explanation

As described above, mixed effects linear models are valuable because they are easily interpreted and allow statements about the statistical significance of predictors. In this section we present another model for the same data which does not offer these benefits but achieves greater predictive accuracy due to its richer modeling framework. This is a *gradient boosting machine* [Chen 2016] built on the same predictors already discussed as well as a number of additional features. These encompass the microbiome (activity of individual organisms and genes), nutrients (weight of meal; subtypes of carbs and fats; micronutrients; specific compounds like caffeine and alcohol), and context (more detailed representations of sleep and activity). Following [Zeevi 2015] we also add two further predictors encoding prior blood glucose levels: the CGM reading immediately before the meal, and the slope of the linear change in CGM readings over the previous 90 minutes. After removing predictors that are low variance, highly correlated with each other, or not correlated with the outcome, a total of 1446 were included in the model.

Data from 82% of users was used to train, with the data from the remaining 18% held out for evaluation. Hyperparameters controlling learning rate, number of trees and tree depth were estimated using cross validation on the training set.

Averaging across 5 such random train/test splits of the data, the model achieves R=.80 (R=.64) on training data, and R=.64 (R=.40) on held-out test data. Performance on training and test data is shown in Figure 4E.

## Discussion

We set out to study the variation of glycemic response based on individual differences, especially differences in gut microbiome activity obtained via the metatranscriptomic method for the first time. We made a few key design choices for the study, including (a) 14 days of monitoring, (b) multiple diet types – omnivore and vegetarian/gluten-free, and (c) a large proportion (66%) of “provided” (pre-designed) meals.

The number of provided meals (104) is considerably larger in our study compared to previous studies (e.g. [Zeevi 2015]; [Mendes-Soares 2019] had only 4 standardized meals). This design allowed us to get more precise readings of consumed meals rather than entirely depending on the smartphone diet tracking app. This choice also ensures that our data allows us to quantify individual PPGR differences between people in response to the same food, reducing the risk that observed differences reflect differences between participants’ diets. Secondly, as shown in Figure 2A, we wanted to design a diet plan that provided broad coverage of the space of macronutrient proportions (carbs, fiber, protein, fat) with the intention of teasing out the independent and interacting effects of the macronutrients. And finally, we needed more control of meals since we also wanted to study the effect of multiple diet types: omnivore and vegetarian/gluten-free (this is ongoing work, not reported here).

It is evident from our data that accounting for individual differences is crucial in providing a full description of PPGRs. We designed 9 specific meals, each of which was a combination of food staples (Figure 2D), that were consumed twice by all participants. While *intra*-person variability for a given meal is substantial (Figure 3C, green), this variability is small relative to the *inter-*person variability for the same meals (blue). We can conclude that while many factors affect PPGRs, some of these factors are individual differences which must be accounted for by differences between people and their lifestyles, not properties of the meal alone. The two meals where intra-person variability is close to inter-person variability are meals that contain little to no carbohydrates.

### Relationships between iAUC and phenotype and food features

We see the expected relationship between age and iAUC in Figure 3E, first panel. The relationship between iAUC and BMI / waist-to-hip ratio (Figure 3E second and third panels) is the opposite of that previously reported in studies such as [Zeevi 2015]. We hypothesize that this may reflect the self-reported good health and high exercise rate within this study population. We see the expected increase in average iAUC with HbA1c in our study population (Figure 3E, fourth panel). However, in our analysis there is no significant effect of HbA1c on iAUC after controlling for other predictors. This may be because the study population was selected for HbA1c in the normal range (< 6.5).

As expected, the bulk of variation in the response is explained by the amount of carbs ingested, and by interactions with fat content in food and other factors that modulate the effect of carbs. Increased fiber resulted in overall lower PPGRs, and while increased fat had little marginal effect by itself, it interacts with carbs to suppress the effect of ingested carbs on the PPGR. The plot of the time-course of this effect in Figure 3B suggests this may happen because fat and protein flatten and delay the digestion of carbs, pushing some of the PPGR out of the 2 hour window considered here ([Franz 1997]; [Wakhloo 1984]). Protein has a numerically negative effect on iAUC but is not significant after controlling for other predictors, so was removed from the final model. Older people had higher PPGRs, and also higher PPGRs per unit of carbs ingested (Figure 4D). Activity after meal consumption as well as sleep immediately before eating both resulted in lower PPGR to carbs, which is consistent with literature on metabolic effects of circadian rhythms [Van Cauter 1997].

The fact that the PPGR is better predicted by the square root of carbs than untransformed carbs is reminiscent of a standard model of gastric emptying in which the volume of food passing from the stomach per unit of time is linear in the square root of its volume [Hopkins 1966].

### Relationships between iAUC and microbiome features

The significant microbiome features related to prediction of postprandial glycemic response were microbiome balance, fucose metabolism pathways, fructose metabolizers, tyrosine metabolizers (marginal), indoleacetate production pathways, and glutamine production pathways (marginal). Of these, fucose, indoleacetate, and tyrosine (marginal) scores interact with carbs, indicating that the microbiome or properties correlated with the microbiome affect the processing of carbs as well as leading to an overall difference in baseline blood sugar.

The microbiome balance score was one of the significant features in predicting glycemic response. Low microbiome balance scores usually result from either an imbalance of relative activities of beneficial vs. harmful microbes or from lower quantity and diversity of microbial organisms. The relationship of suboptimal overall gut microbiome and higher PPGR is in line with the current literature [Karlsson 2013; Larsen 2010; Vrieze 2012] implicating the role of gut health in glycemic regulation.

The fucose metabolism pathway score showed a direct relationship with PPGR. Fucose is a sugar molecule that various microbial organisms can use as an energy source [Chen 1987]. When other carbohydrate sources are not available, gut microbiota can switch to using the fucose that can be obtained from the host’s gut mucosal lining. This process is often carried out by microbes known as mucin degraders, such as certain species of genus Ruminococcus [Crost 2016]. We therefore hypothesize that higher fucose consumption activity, as reflected by the microbiome pathway score, may be associated with microbiomes of those individuals who are either more likely to fast or whose internal ecosystem and overall body state resembles the conditions of fasting or calorie deprivation. This may explain its association with higher PPGR. More research is needed to establish relationships between microbial metabolism of gut sugars and the host’s tendency to show higher glucose spikes in the blood after meals.

The indoleacetate production pathways score incorporates the role and significance of expressed genes in the context of microbial indoleacetate production. The algorithm takes all the known pathway axes that ultimately lead to microbial production of compounds of type indole acetic acids (IAAs) and scores them using gene expression as input data. In the case of indole acetate production, the result shows that when such pathways score “High” in activity, the glycemic response to the given food is lower. This is consistent with known anti-inflammatory properties of IAA [Krishnan 2018; Whitfield-Cargile 2016]. Inflammatory activities in the gut and their consequential potential to cause systemic low-grade inflammation are implicated in the development of Type 2 Diabetes and other metabolic disorders [Gonzalez 2018; Tuomainen 2018]. Moreover, there are direct implications of IAAs in glycemic response, and some findings suggest hypoglycemic action of indole-3-acetic acid in diabetes mellitus [Mirsky 1956].

Indoles and indoleacetate are beneficial products of protein fermentation, and tryptophan metabolism pathway products particularly [Russell 2013]. In one study, intraperitoneal administration of indole-3-propionic acid, indole-3-butyric acid, and indole-3-acetic acid were shown to be associated with hypogIycemia in normal and alloxandiabetic mice, while L-tryptophan and kynurenic acid had no effect [Silverstein 1966].

The interpretation we offer here is dependent on the individual’s microbiome function. If a given person’s microbiota mainly shows ability to convert tryptophan to beneficial indoles and indole-actetate molecules capable of reducing inflammation and glycemic effects of foods, then it may be of benefit to recommend tryptophan sources (in the form of food or supplement) to such people. On the other hand, if tryptophan is used by the microbiota to produce more of the pro-inflammatory triggers, then such action may not be suitable for mitigating glycemic response or inflammation in general.

Tyrosine and fructose metabolizer scores group active microbes by functional characteristics. In our studies we have observed that active functional microbial groups reflect the host’s habitual diet. Hence, “Low” tyrosine metabolizers may suggest a diet that is low on protein sources of tyrosine. We also hypothesize that the inverse relationship between the fructose metabolizers score and PPGR may be due to a diet that is low in fructose, or other carbohydrates that serve as metabolic precursors of fructose. It is not yet clear how a diet that is rich in fructose or deficient in tyrosine may influence glycemic response, and more studies are needed.

Glutamine production pathways score, when “Low,” showed a direct relationship with higher PPGR. Microbial glutamine production has not been directly linked to glycemic response in humans. However, glutamine is considered an important nutrient for gut health and has been included in various supplements used by clinical healthcare practitioners to prevent or heal “leaky gut” [Kim 2017, Rao 2012]. More research is needed to understand the molecular mechanisms that may be responsible for higher glycemic response to food in individuals with low microbial glutamine production activity in the gut.

The microbiome features revealed by our glycemic response model may influence PPGR directly or indirectly. Although it is challenging to delineate causal mechanisms, there may be functional patterns that connect the significant scores with gut health, intestinal barrier integrity, and inflammation. Inflammation and stress response may be implicated in elevation of blood glucose (either due to cortisol pathway or other mechanisms). Knowing which foods may elicit higher personal PPGR can offer valuable guidance in diet selection. However, to intervene on the root cause of glycemic response, specific mechanisms connecting nutrients to the gut microbiome and to inflammatory and glycemic response need to be taken into account. We seek to confirm and validate these mechanisms. An understanding of which microbiome features are significant will pave the path to precise personalization of food and supplement recommendations.

### Modeling methods and model evolution

The multilevel mixed effects model presented first in the previous section was deliberately chosen to better understand the incremental effects of the significant features, especially the functional gut microbiome activity features. We are not aware of any prior literature that demonstrates the statistical significance of the microbiome in the context of a predictive model of PPGR. Prior studies [Zeevi 2015] using only ensemble methods such as gradient boosting machines represent the state of the art in accurate prediction of PPGR (which we also show in Figure 4E). These models suffer from difficulty of interpretation, including determination of which features significantly contribute to the predicted outcome.

We are in the process of increasing the generalizability of our findings by collecting further data from underrepresented subpopulations such as pre-diabetics, people reporting poor overall health, and older participants. With the goal of continuous improvement, we will rebuild and revalidate our model based on this new expanded data. The current paper provides a first snapshot of the collected data, and we will use the additional data to consolidate the current model, as well as potentially surface new relevant predictors. Finally, we also plan on validating the model on an unseen cohort, and performing a blinded randomized controlled dietary intervention based on this predictive model to look for improvements in the glycemic response as well as alterations to the gut microbiota.

## Conclusions

Most significantly, this paper makes the following contributions:

- Demonstrates for the first time that *metatranscriptomic* activity of the gut microbiome contributes to individual variation in glycemic response among adults.
- Suggests new microbial features that may help uncover molecular mechanisms of glycemic control.
- Demonstrates the statistical significance of all features using a multilevel mixed-effects regression model where fixed effects represent measured properties and random effects account for further variation. We also present a gradient boosting machine for maximizing predictive accuracy.
- Demonstrates that glycemic response is driven by the properties of an individual in addition to the food’s macronutrient content, measured with 104 unique pre-designed meals within omnivore and vegetarian/gluten-free diet types and within a multi-ethnic population.

## Acknowledgements

We are grateful to the Viome study team – (alphabetically) Ashleigh Winter, Barry Zhao, Braidon Waggoner, Brendan Zapp, Calita Quesada, Erica Castillo, Gabby Hands, James Horne, Kelly Nebgen, Marcela Walker, Miranda Intrator, Pete Lichtman, Ryan Toma, Sabria Hankins, Tristen Hill, and YehShen McShan – for handling the considerable logistics during the data collection and annotation phases. We thank Matteo Rinaldi and Sha Yu Han for initial data exploration and analyses; Subha Krishnan for curating the microbiome pathway literature and features; and Debra Heald and Samika Savanur for their clinical input.

## Author contributions

G.B., H.M., and M.V. designed the study. A.S. managed the study logistics including sample and clínical data collection. M.V. performed the lab analysis. H.L. developed the meal plans, and E.P. organized and analyzed the nutrient data, and contributed to writing. N.K. developed the bioinformatics pipeline. H.T., S.G., M.G., V.G., and I.S. performed the analyses, developed the machine learning models, generated visualizations, and contributed to writing. A.P. and H.M. interpreted the features and contributed to writing. H.T. and G.B. guided the analysis and led the manuscript writing.

## References

https://www.accessdata.fda.gov/cdrh_docs/pdf15/p150021c.p df

Bashiardes S., Zilberman-Schapira G., Elinav E. 2016. Use of Metatranscriptomics in Microbiome Research. Bioinformatics and Biology Insights 10:BBI.S34610. DOI: 10.4137/BBI.S34610.

Bates, Douglas, Martin Maechler, Ben Bolker, Steven Walker, Rune Haubo Bojesen Christensen, Henrik Singmann, Bin Dai, Gabor Grothendieck, C+ Eigen, and L. Rcpp. “Package ‘lme4’.” Convergence 12, no. 1 (2015).

Indra Bervoets, Daniel Charlier, Diversity, versatility and complexity of bacterial gene regulation mechanisms: opportunities and drawbacks for applications in synthetic biology, FEMS Microbiology Reviews, Feb 2019, https://doi.org/10.1093/femsre/fuz001

https://www.cdc.gov/media/releases/2017/p0718-diabetes-report.html

Chen, Y. M., Zhu, Y., Lin, E. C. “NAD-linked aldehyde dehydrogenase for aerobic utilization of L-fucose and L-rhamnose by Escherichia coli.” J Bacteriol. 1987;169(7):3289–94

Chen, T., Guestrin, C. “XGBoost: A scalable tree boosting system” KDD’16, Proceedings of the 22nd ACM SIGKDD International Conference on Knowledge Discovery and Data Mining (2016): 785–794.

Cheng, J. T. “Merit of Incremental Area under the Curve (iAUC) in Nutrition is Varied in Pharmacological Assay-A Review.” Clin J Dia Care Control 1, no. 2 (2018): 180008.

Crost EH, Tailford LE, Monestier M, Swarbreck D, Henrissat B, Crossman LC, Juge N.” The mucin-degradation strategy of Ruminococcus gnavus: The importance of intramolecular trans-sialidases” Gut Microbes. 2016 Jul 3;7(4):302–312.

Franz, Marion J. “Protein: metabolism and effect on blood glucose levels.” The diabetes educator 23, no. 6 (1997): 643–651.

Gelman, Andrew, and Jennifer Hill. “Data analysis using regression and hierarchical/multilevel models.” New York, NY: Cambridge (2007).

Lopez, Laura, Karin Garrie, and Mark D. Turner. “Type 2 diabetes–An autoinflammatory disease driven by metabolic stress.” Biochimica et Biophysica Acta (BBA)-Molecular Basis of Disease (2018).

Gosalbes MJ., Durbán A., Pignatelli M., Abellan JJ., Jiménez-Hernández N., Pérez-Cobas AE., Latorre A., Moya A. 2011. Metatranscriptomic Approach to Analyze the Functional Human Gut Microbiota. PLoS ONE 6:e17447. DOI: 10.1371/journal.pone.0017447.

Gutierrez, Mario, Mehras Akhavan, Lois Jovanovic, and Charles M. Peterson. Utility of a short-term 25% carbohydrate diet on improving glycemic control in type 2 diabetes mellitus. Journal of the American College of Nutrition 17.6 (1998): 595–600. DOI: 10.1080/07315724.1998.10718808

Andrew Hatch, James Horne, Ryan Toma, Brittany L Twibell, Kalie M Somerville, Benjamin Pelle, Kinga P Canfield, Guruduth Banavar, Ally Perlina, Helen Messier, Niels Klitgord, and Momchilo Vuyisich. 2019. A robust metatranscriptomic technology for population-scale studies of diet, gut microbiome, and human health. doi:10.31219/osf.io/8vd6x

He S., Wurtzel O., Singh K., Froula JL., Yilmaz S., Tringe SG., Wang Z., Chen F., Lindquist EA., Sorek R., Hugenholtz P. 2010. Validation of two ribosomal RNA removal methods for microbial metatranscriptomics. Nature Methods 7:807–812. DOI: 10.1038/nmeth.1507.

Hopkins A: The pattern of gastric emptying: a new view of old results. J Physiol (Lond) 182:144-149, 1966

Jenkins David JA, and Alexandra L. Jenkins. “Dietary fiber and the glycemic response.” Proceedings of the Society for Experimental Biology and Medicine 180, no. 3 (1985): 422–431. DOI: 10.3181/00379727-180-42199

Karlsson, F. H., Tremaroli, V., Nookaew, I., Bergström, G., Behre, C. J., Fagerberg, B., Nielsen, J., Bäckhed, F. “Gut metagenome in European women with normal, impaired and diabetic glucose control.” Nature (2013) 498(7452):99–103.

KEGG: Kyoto Encyclopedia of Genes and Genomes. http://www.kegg.jp/ or http://www.genome.jp/kegg/

Min-Hyun Kim and Hyeyoung Kim. “The Roles of Glutamine in the Intestine and Its Implication in Intestinal Diseases” Int J Mol Sci. (2017) 18(5): 1051.

Knight R, Vrbanac A, Taylor BC, Aksenov A, Callewaert C, Debelius J, Gonzalez A, Kosciolek T, McCall L-I, McDonald D, Melnik AV, Morton JT, Navas J, Quinn RA, Sanders JG, Swafford AD, Thompson LR, Tripathi A, Xu ZZ, Zaneveld JR, Zhu Q, Caporaso JG, Dorrestein PC. 2018. Best practices for analysing microbiomes. Nature Reviews Microbiology 16:410–422. DOI: 10.1038/s41579-018-0029-9.

Krishnan S, Ding Y, Saedi N, Choi M, Sridharan GV, Sherr DH, Yarmush ML, Alaniz RC, Jayaraman A, Lee K. ” Derived Tryptophan Metabolites Modulate Inflammatory Response in Hepatocytes and Macrophages” Cell Rep. 2018 Apr 24;23(4):1099–1111

Larsen, N., Vogensen, F. K., van den Berg, F. W., Nielsen, D. S., Andreasen, A.S., Pedersen, B.K., Al-Soud, W. A., Sørensen, S.J., Hansen, L.H., Jakobsen, M. “Gut microbiota in human adults with type 2 diabetes differs from non-diabetic adults.” PLoS One. 2010;5(2):e9085

Livesey, Geoffrey, Richard Taylor, Toine Hulshof, and John Howlett. “Glycemic response and health—a systematic review and meta-analysis: relations between dietary glycemic properties and health outcomes.” The American journal of clinical nutrition 87, no. 1 (2008): 258S–268S. DOI: 10.1093/ajcn/87.1.258S

Li J, Jia H, Cai X, Zhong H, Feng Q, Sunagawa S, Arumugam M, Kultima JR, Prifti E, Nielsen T, Juncker AS, Manichanh C, Chen B, Zhang W, Levenez F, Wang J, Xu X, Xiao L, Liang S, Zhang D, Zhang Z, Chen W, Zhao H, Al-Aama JY, Edris S, Yang H, Wang J, Hansen T, Nielsen HB, Brunak S, Kristiansen K, Guarner F, Pedersen O, Doré J, Ehrlich SD, Bork P, Wang J. 2014. “An integrated catalog of reference genes in the human gut microbiome.” Nature Biotechnology 32:834–841. (2014) DOI: 10.1038/nbt.2942.

Ludwig David S., Frank B. Hu, Luc Tappy, and Jennie Brand-Miller. “Dietary carbohydrates: Role of quality and quantity in chronic disease.” Bmj 361 (2018): k2340. DOI: 10.1136/bmj.k2340

Mendes-Soares H, Raveh-Sadka T, Azulay S, et al. Assessment of a Personalized Approach to Predicting Postprandial Glycemic Responses to Food Among Individuals Without Diabetes. JAMA Network Open. 2019;2(2):e188102. doi:10.1001/jamanetworkopen.2018.8102

Mirsky, I. Arthur, and Daniel Diengott. “Hypoglycemic action of indole-3-acetic acid by mouth in patients with diabetes mellitus.” Proceedings of the Society for Experimental Biology and Medicine 93, no. 1 (1956): 109-110.

[Perlina in prep.]Perlina et al, “Pathway Scoring and Functional Assessment of Microbiome Activities Connects Gut Metatranscriptomics to Phenotype and Health Metrics.” -in preparation

RadhaKrishna Rao and Geetha Samak “Role of Glutamine in Protection of Intestinal Epithelial Tight Junctions” J Epithel Biol Pharmacol. 2012 Jan; 5(Suppl 1-M7): 47–54.

Russell Wendy R., Sylvia H. Duncan, Lorraine Scobbie, Gary Duncan, Louise Cantlay, A. Graham Calder, Susan E. Anderson, and Harry J. Flint. “Major phenylpropanoid-derived metabolites in the human gut can arise from microbial fermentation of protein.” Molecular nutrition & food research 57, no. 3 (2013): 523-535.

Silverstein Murray N., Khalil G. Wakim, Robert C. Bahn, and Richard H. Decker. “Role of tryptophan metabolites in hypoglycemia associated with neoplasia.” Cancer 19, no. 1 (1966): 127–133.

Suez J, Shapiro H, Elinav E. Role of the microbiome in the normal and aberrant glycemic response. Clin Nutr Exp. 2016;6:59–73. doi:10.1016/j.yclnex.2016.01.001

Tuomainen M, Lindström J, Lehtonen M, Auriola S, Pihlajamäki J, Peltonen M, Tuomilehto J, Uusitupa M, Mello V and Hanhineva K. “Associations of serum indolepropionic acid, a gut microbiota metabolite, with type 2 diabetes and low-grade inflammation in high-risk individuals” Nutrition & Diabetes 35 (2018)

Van Cauter, Eve, Kenneth S. Polonsky, and André J. Scheen. “Roles of circadian rhythmicity and sleep in human glucose regulation.” Endocrine reviews 18, no. 5 (1997): 716-738.

Vrieze, A., Van Nood, E., Holleman, F., Salojärvi, J., Kootte, R. S., Bartelsman, J. F., Dallinga-Thie, G. M., Ackermans, M. T., Serlie, M. J., Oozeer, R., Derrien, M., Druesne, A., Van Hylckama Vlieg, J. E., Bloks, V. W., Groen, A.K., Heilig, H. G., Zoetendal, E. G., Stroes, E. S., de Vos, W. M., Hoekstra, J. B., Nieuwdorp, M. “Transfer of intestinal microbiota from lean donors increases insulin sensitivity in individuals with metabolic syndrome.” Gastroenterology. 2012 Oct;143(4):913-6.e7

Wakhloo, A. K., J. Beyer, C. Diederich, and G. Schulz. “Effect of dietary fat on blood sugar levels and insulin consumption after intake of various carbohydrate carriers in type I diabetics on the artificial pancreas.” Deutsche medizinische Wochenschrift (1946) 109, no. 42 (1984): 1589-1594.

Whitfield-Cargile Canaan M., Noah D. Cohen, Robert S. Chapkin, Brad R. Weeks, Laurie A. Davidson, Jennifer S. Goldsby, Carrie L. Hunt et al. “The microbiota-derived metabolite indole decreases mucosal inflammation and injury in a murine model of NSAID enteropathy.” Gut microbes 7, no. 3 (2016): 246-261.

Wolever, T.M., and Jenkins, D.J. (1986). The use of the glycemic index in predicting the blood glucose response to mixed meals. Am. J. Clin. Nutr. 43, 167–172.

David Zeevi, Tal Korem, Niv Zmora, David Israeli, Daphna Rothschild, Adina Weinberger, Orly Ben-Yacov, Dar Lador, Tali Avnit-Sagi, Maya Lotan-Pompan, et al. Personalized Nutrition by Prediction of Glycemic Responses.Cell. 2015 Nov 19; 163(5): 1079–1094. doi: 10.1016/j.cell.2015.11.001

